# Island biogeography theory and the gut: why taller people tend to harbor more diverse gut microbiomes

**DOI:** 10.1101/2023.08.08.552554

**Authors:** Katherine Ramos Sarmiento, Alex Carr, Christian Diener, Kenneth J. Locey, Sean M. Gibbons

## Abstract

Prior work has shown a positive scaling relationship between vertebrate body size and gut microbiome alpha-diversity. This observation mirrors commonly observed species area relationships (SAR) in many other ecosystems. Here, we show a similar scaling relationship between human height and gut microbiome alpha-diversity across two large, independent cohorts, controlling for a wide range of relevant covariates, such as body mass index, age, sex, and bowel movement frequency. Island Biogeography Theory (IBT), which predicts that larger islands tend to harbor greater species diversity through neutral demographic processes, provides a simple mechanism for these positive SARs. Using an individual-based model of IBT adapted to the gut, we demonstrate that increasing the length of a flow-through ecosystem is associated with increased species diversity. We delve into the possible clinical implications of these SARs in the American Gut Cohort. Consistent with prior observations that lower alpha-diversity is a risk factor for *Clostridioides difficile* infection (CDI), we found that individuals who reported a history of CDI were shorter than those who did not and that this relationship appeared to be mediated by alpha-diversity. We also observed that vegetable consumption mitigated this risk increase, also by mediation through alpha-diversity. In summary, we find that body size and gut microbiome diversity show a robust positive association, that this macroecological scaling relationship is related to CDI risk, and that greater vegetable intake can mitigate this effect.

## Introduction

From the moment we are born, we are colonized by a diverse community of commensal microbiota that we carry with us throughout our lives ^1^. The vast majority of our commensal microbes reside in the gastrointestinal tract ^2^. Our gut microbiota has an enormous impact on our phenotype ^3^, with almost half of the metabolites circulating in human blood significantly associated with cross-sectional variation in the ecological composition of the gut microbiome ^4,5^. One of the key ecosystem functions that the gut microbiota provides to its host is resistance to enteric bacterial pathogens ^6^. Niche saturation or nutrient competition are commonly invoked mechanisms for how the microbiota excludes invaders ^6^. Specifically, species-diverse commensal communities are more apt to saturate available metabolic niches so that an invasive pathogen is less likely to colonize, out-compete commensals, and cause disease ^6,7^. Though many determinants of gut microbiome alpha-diversity (i.e., taxon richness and/or evenness in a given sample) are known, including diet, intestinal transit time, and antibiotic treatment, much of the variation in gut alpha-diversity remains unexplained ^8–11^.

Vertebrate body size, which varies over six orders of magnitude, has been shown to be positively associated with gut microbiome alpha-diversity, indicating that larger animals with larger guts harbor more species ^12^. This pattern mirrors similar species area relationships (SARs) seen in other ecosystems, where larger areas correspond to more observed taxa ^13–15^. Similarly, recent work from a large human cohort showed a positive association between height, which only varies by approximately two-fold across people, and gut microbiome alpha-diversity^16^. We expanded on this observation across two independent human cohorts (i.e., the Arivale and American Gut cohorts), and found that the association between human height and gut alpha-diversity was highly robust to the inclusion of several covariates known to influence alpha-diversity, like body mass index (BMI), bowel movement frequency (BMF), age, and sex. The mechanisms underlying these gut microbiome SARs, and their potential clinical consequences, have yet to be explored.

Island Biogeography Theory (IBT), a classic neutral model of species immigration/emigration, birth/death, and speciation/extinction, predicts a positive SAR ^17^. Specifically, larger islands tend to harbor more individuals, which ultimately gives rise to a larger number of coexisting species ^17^. We hypothesized that this simple neutral model may explain the size-diversity relationships observed across vertebrates and within humans. To further explore this hypothesis, we built an individual-based model (IBM) that approximates IBT in the gut, allowing for variation in system length, with immigration, emigration, birth, death, and a unidirectional flow through the system. We simulated length ranges that approximate the scaling of vertebrate body sizes and length ranges that approximate human height variation, to compare these results to our empirical observations. Overall, we find IBT to be a credible mechanism for the rather noisy, but statistically robust relationship we observe between human height and gut microbiome alpha-diversity.

Finally, we explored the potential clinical implications of this scaling between height and gut microbiome alpha-diversity. Lower gut microbiome alpha-diversity has been associated with greater susceptibility to enteric infections ^7,18^. By extension of the scaling between height and gut microbiome alpha-diversity, individuals who suffer from a greater frequency of enteric infections may tend to be shorter than average. Indeed, we found that participants in the American Gut cohort who reported a history of *Clostridioides difficile* infection (CDI) were significantly shorter and had less diverse gut microbiomes than those who did not report a history of CDI. In addition, we leverage the American Gut dietary metadata to show that higher vegetable consumption, which is known to increase gut alpha-diversity ^8^, can modify this association between height, diversity, and CDI. Finally, we found that gut alpha-diversity was a significant mediator of the effects of both height and dietary intake on CDI history, which is consistent with the niche saturation hypothesis that posits that a more diverse commensal microbiota protects the host from enteric pathogens through competitive exclusion ^19,20^.

## Results

### Positive scaling between vertebrate body size and gut microbiome alpha-diversity

Godon et al. found that vertebrate body size and gut microbiome alpha-diversity were positively associated ^12^. This study used an older method for quantifying alpha-diversity that involves rapidly transitioning ssDNA (16S amplicons) from warm to cold temperatures so that strands fold into unique shapes that are determined by their primary sequence. These unique, negatively-charged ssDNA strands will migrate at different rates through a capillary tube during electrophoresis, and the differing peaks of fluorescence are used to calculate taxon diversity ^12,21^. We also leveraged 16S amplicon sequencing data from vertebrate guts, spanning a wide range of body sizes ^22,23^. We re-analyzed the Groussin et al. (n=32) and Song et al. (n=1,373) data sets (**Fig. 1A**), along with the Godon et al. data set (n=80, **Fig. 1B**), and observed a consistent, positive association between log-Simpson’s Diversity and log-mass across all three studies (**Fig. 1**). Ordinary Least Squares (OLS) regression analysis (log-Simpson’s Diversity ∼ log-mass) showed similar variances explained across data sets (*r*^2^∼ 0.5, *p* < 1.83 · 10^-14^).

**Figure 1.**
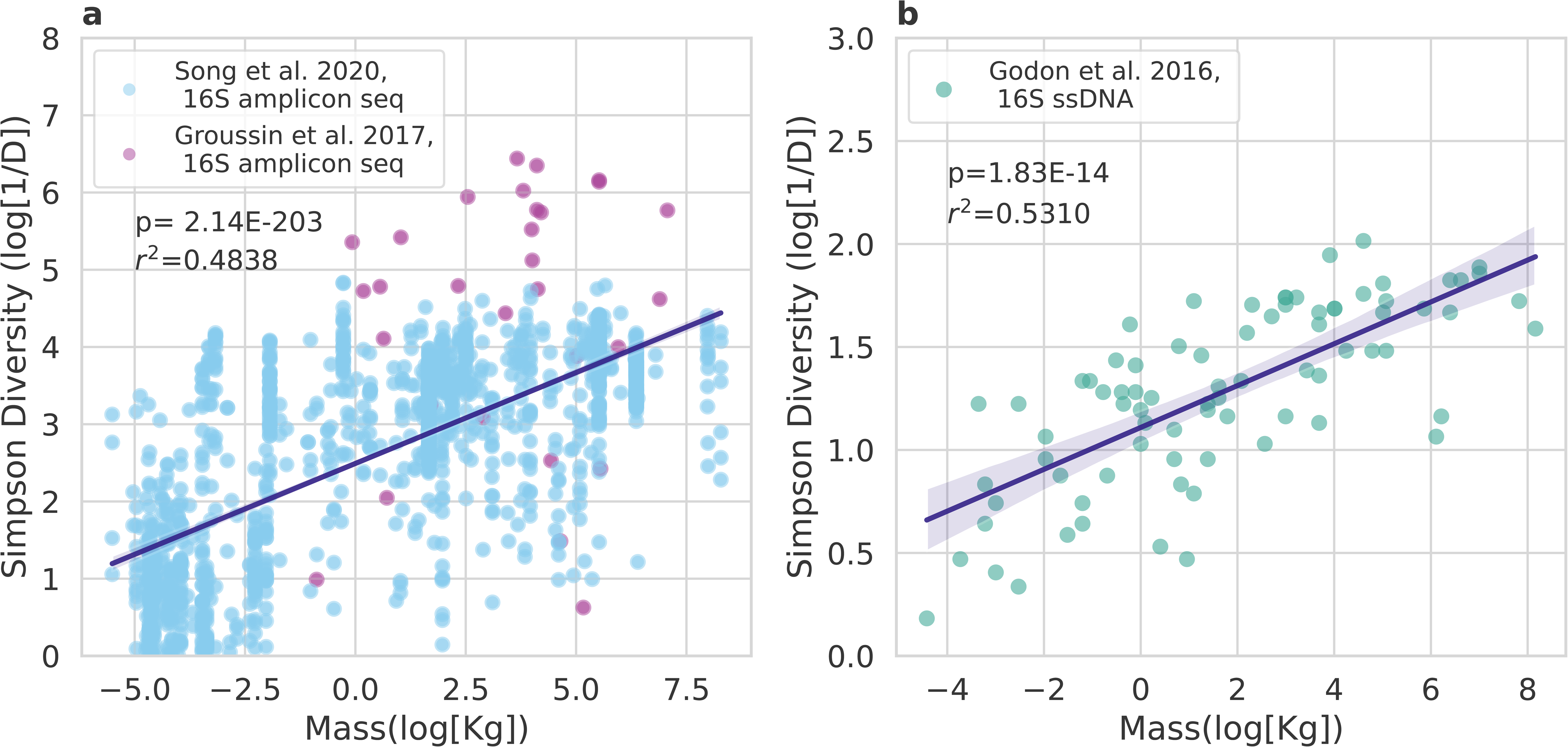
Relationship between body size and gut microbiome Simpson’s Diversity across vertebrates. (A) Log vertebrate body mass and log gut microbiome Simpson’s Diversity are positively associated across two independent 16S amplicon sequencing data sets (Song et al. and Groussin et al.). Ordinary Least Squares regression analysis shows this association is statistically significant (*r*^2^ = 0.4838, *p* < 10^-16^). (B) A similar result emerges from a CE-SSCP data set (*r*^2^ = 0.5310, *p* = 1.83 * 10^-14^).

### Relationship between human height and gut microbiome alpha-diversity is robust to inclusion of relevant covariates

We found that human height and gut microbiome alpha-diversity were positively associated across two large human cohorts. We used data from the Arivale cohort (n=3,067) and the American Gut cohort (n=5,572) to compare log-Simpson’s Diversity versus log-height (**Fig. 2**). The Arivale cohort consisted of self-selected American adults who had enrolled in a scientific wellness program, and the American Gut cohort consisted of self-selected adults participating in a citizen-science program, primarily from the United States, the United Kingdom, and Australia^8,24^.

**Figure 2.**
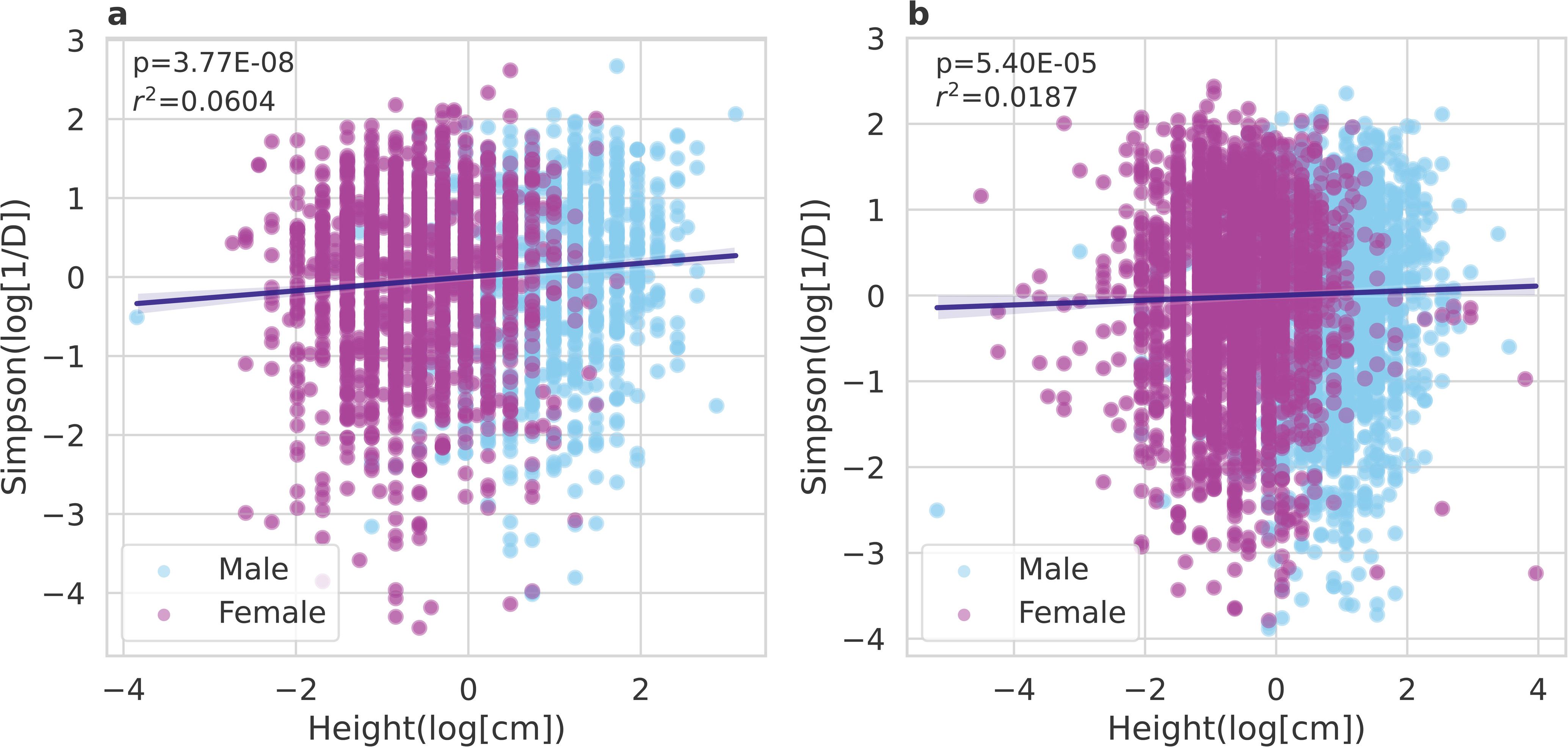
Relationship between height and gut microbiome Simpson’s Diversity across humans. Both figures displayed are log-log regression plots displaying Simpson’s Diversity versus height in (A) the Arivale cohort (n=3,067) and (B) the American Gut cohort (n=5,572). Both plots show a similar trend where log-height is positively associated with log-Simpson’s Diversity. Ordinary Least Squares regressions used the formula: log-Simpson’s Diversity∼ sex + age + BMI + BMF + log-mass. OLS on the Arivale cohort yielded a model *r*^2^ = 0.06 and a *p* = 3.77 * 10^-8^. OLS on the American Gut cohort yielded a *r*^2^ = 0.0187 and a *p* = 5.40 * 10^-5^.

Because several demographic variables are known to affect gut microbiome diversity, we ran OLS regression on each cohort individually, with age, sex, body mass index (BMI), and bowel movement frequency (BMF) variables as covariates. Consistent with the literature, we found that in both cohorts, being male, having a higher BMF, and a higher BMI were all negatively associated with log-Simpson’s Diversity, and age was positively associated with log-Simpson’s Diversity (**Fig. 2**). Not only was log-height positively associated with Simpson’s Diversity in the absence of covariates, but it retains its significance in the presence of these covariates (**Table 1**). Moreover, performing an ANOVA F-test, comparing a reduced model without height versus a full model including height, yielded a significantly higher fraction of explained variability across both cohorts (*p* = 3.77 · 10^-8^; **Table 1**).

**Table 1:** Ordinary Least Squares (OLS) Regression shows height’s association with Simpson’s Diversity is robust to confounders. Height retains its signal with and without the inclusion of covariates across both cohorts. When we include covariates, height has a *r*^2^ = 0.0093 and *p* = 3.77 • 10^-8^ in the Arivale cohort, and a *r*^2^ = 0.0029 and *p* = 5.4 • 10^-5^ in the American Gut cohort. When we exclude covariates, the Arivale cohort yields a model *r*^2^ = 0.0076 and *p* = 1.28 • 10^-6^ and the American Gut cohort yields a model *r*^2^ = 0.0008 and *p* = 0.040. Moreover, F-testing of the model comparing a reduced model using only the covariates to a full model including the covariates and height yields a significant p-value (*p* = 3.77 • 10^-8^ for the Arivale cohort, and *p* = 5.40 • 10^-5^ for the American Gut cohort).

| cohort | covariate | $r^2$ (covariate) | $p$ (covariate) | Beta | formula | $r^2$ | F | p (F) |
| --- | --- | --- | --- | --- | --- | --- | --- | --- |
| Arivale | sex[T.M] | 0.0016 | 2.15E-02 | -0.1180 | simpson~age+sex+BMI+BMF+ height | 0.0604 | 30.42 | 3.77E-08 |
|  | age | 0.0059 | 1.31E-05 | 0.0769 | simpson~age+sex+BMI+BMF+ height | 0.0604 | 30.42 | 3.77E-08 |
|  | BMI | 0.0338 | 2.32E-25 | -0.1847 | simpson~age+sex+BMI+BMF+ height | 0.0604 | 30.42 | 3.77E-08 |
|  | BMF | 0.0088 | 9.63E-08 | -0.0958 | simpson~age+sex+BMI+BMF+ height | 0.0604 | 30.42 | 3.77E-08 |
|  | height | 0.0093 | 3.77E-08 | 0.1368 | simpson~age+sex+BMI+BMF+ height | 0.0604 | 30.42 | 3.77E-08 |
|  | height |  | 1.28E-06 | 0.0873 | simpson~height | 0.0076 | 23.55 | 1.28E-06 |
| American Gut | sex[T.M] | 0.0014 | 5.56E-03 | -0.1025 | simpson~age+sex+BMI+BMF+ height | 0.0187 | 16.33 | 5.40E-05 |
|  | age | 0.0084 | 5.64E-12 | 0.0928 | simpson~age+sex+BMI+BMF+ height | 0.0187 | 16.33 | 5.40E-05 |
|  | BMI | 0.0011 | 1.26E-02 | -0.0340 | simpson~age+sex+BMI+BMF+ height | 0.0187 | 16.33 | 5.40E-05 |
|  | BMF | 0.0070 | 3.29E-10 | -0.0842 | simpson~age+sex+BMI+BMF+ height | 0.0187 | 16.33 | 5.40E-05 |
|  | height | 0.0029 | 5.40E-05 | 0.0736 | simpson~age+sex+BMI+BMF+ height | 0.0187 | 16.33 | 5.40E-05 |
|  | height |  | 4.02E-02 | 0.0275 | simpson~height | 0.0008 | 4.21 | 4.02E-02 |

### Adapting Island Biogeography Theory to the gut

In order to demonstrate a mechanistic link between body size and gut alpha-diversity, we formulated IBT in a simplified model designed to simulate varying gut lengths (**Fig. 3**).

**Figure 3.**
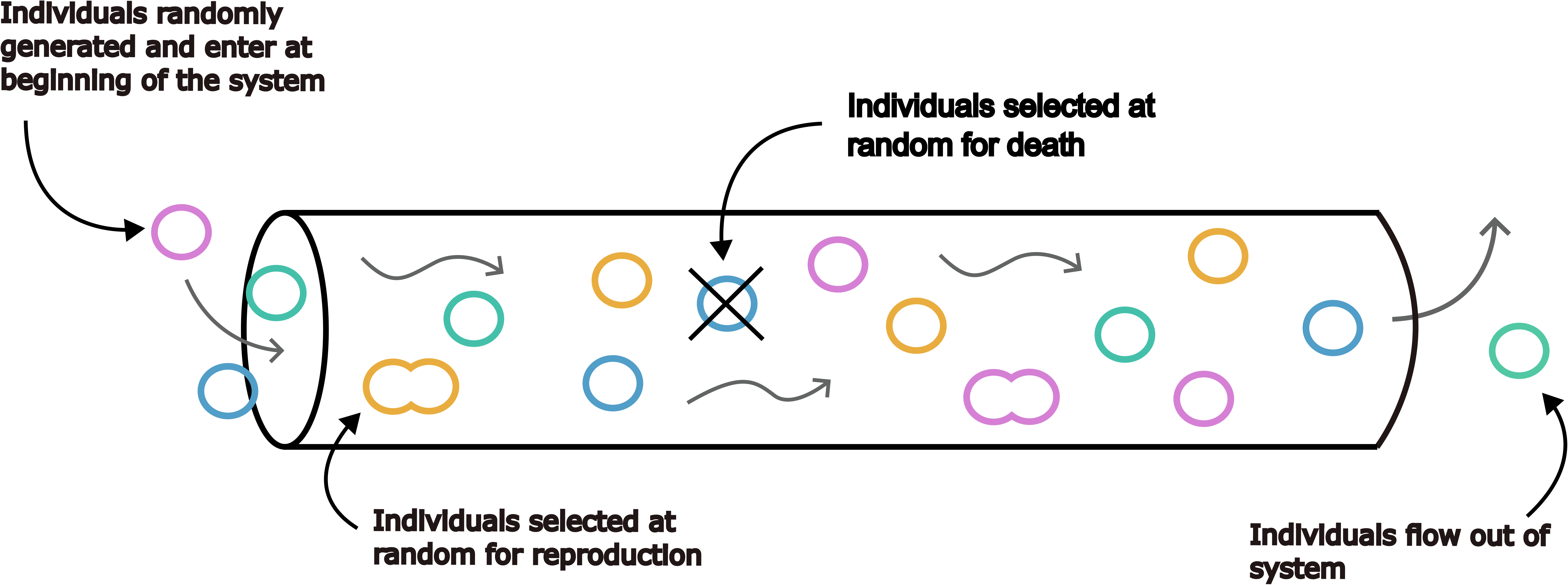
A schematic of the Individual-Based Model (IBM) used to simulate Island Biogeography Theory (IBT) in the gut. We built a simple IBM that approximated the unidirectional flow of the gut, where we could vary the length of the system. Individuals were randomly generated from a heavy-tailed species abundance distribution, entering the system on the left side, flowing along the length of the system over time, and eventually exiting the right side of the system. The number of individuals entering the system per time step were determined by the immigration rate of the simulation. In addition to flowing through the system, some individuals were randomly selected for reproduction or death at each time step.

We ran 2 sets of 1000 simulations, one approximating the vertebrate size range (six orders of magnitude; **Fig 4A**) and another approximating the human size range (∼2X; **Fig 4B**). In both sets of simulations, Simpson’s Diversity was positively associated with simulated intestine length (**Fig. 4**). OLS regression (log[Simpson (1/D)] ∼ log[system length]) showed that the simulations run on the vertebrate scale had a much larger *r*^2^ value (*r*^2^ = 0.3404, *p* = 2.96 •10^-92^) compared to the simulations run on the human scale (*r*^2^ = 0.0703, *p* = 1.50 • 10^-17^).

**Figure 4.**
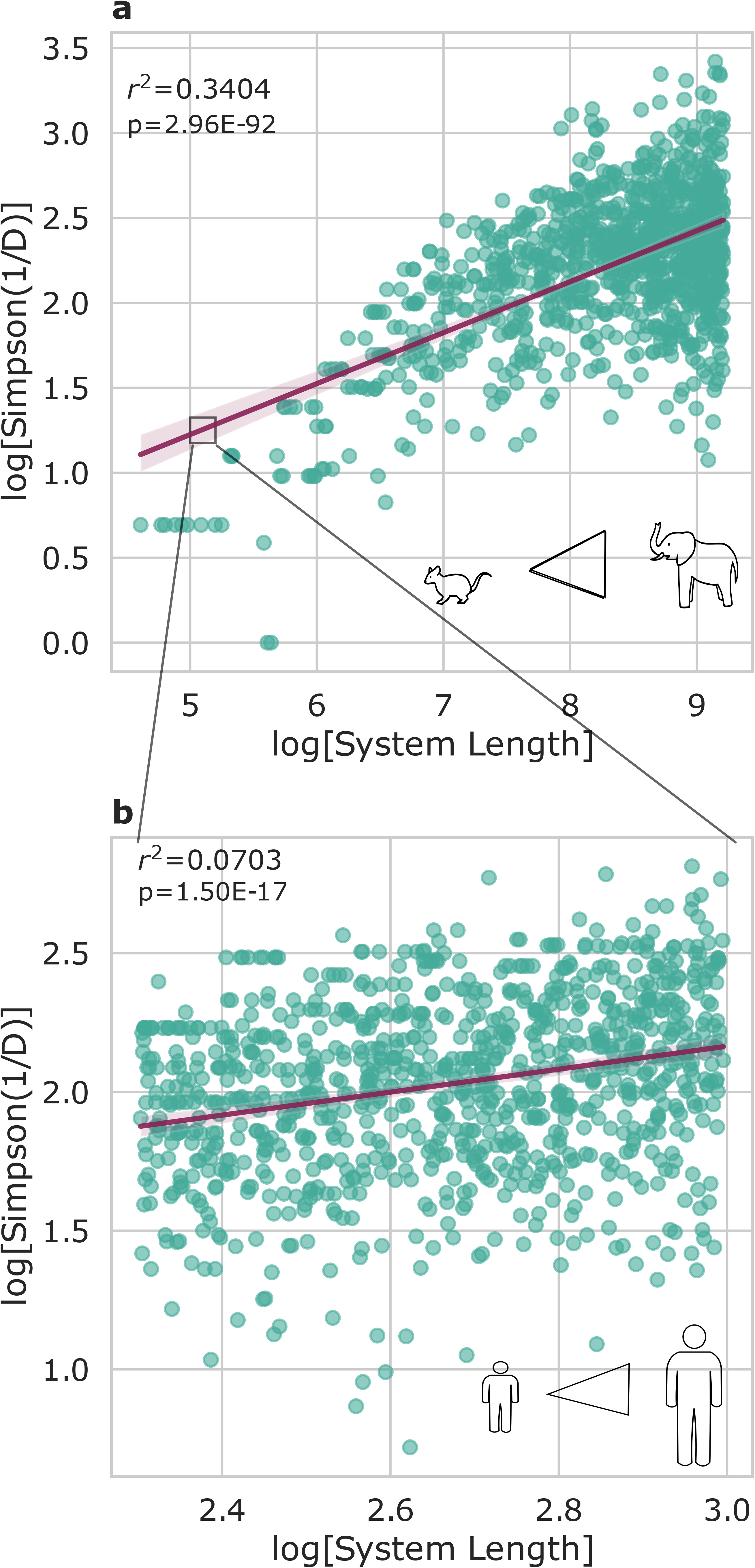
Simulations mirror empirical relationships observed between size and diversity. Both plots show 1000 simulations, where the length of the system was varied within a set range. (A) When varying IBM lengths over six orders of magnitude, similar to the size range we observed across vertebrates, we see a strong association between size and diversity (OLS *r*^2^ = 0.3404 and *p* = 2.96 * 10^-92^). (B) When varying IBM length over a much smaller two-fold range, which is in line with the observed range in human heights, we see a much weaker association between size and diversity, similar to empirical observations (OLS *r*^2^ = 0.0703 and *p* = 1.50 * 10^-17^). The box drawn in this figure is not to scale but has been increased in size for ease of visibility.

### Exploring the clinical implications of the height-diversity relationship in the American Gut cohort

Low gut alpha-diversity has been associated with susceptibility to enteric infections ^7^. Therefore, we asked whether or not shorter individuals, with slightly lower alpha-diversity, might be more susceptible to CDI. The American Gut cohort contained individuals with self-reported histories of CDI, which allowed us to explore this hypothesis.

Consistent with prior literature, individuals who reported a history of CDI had less diverse gut microbiomes than those who did not (Welch’s t-test t=4.92, *p* ≤ 0.0001; **Fig 5A**). Moreover, individuals who reported a history of CDI were shorter than those who did not (Welch’s t-test t=3.92, *p* ≤ 0.001; **Fig. 5D**).

**Figure 5.**
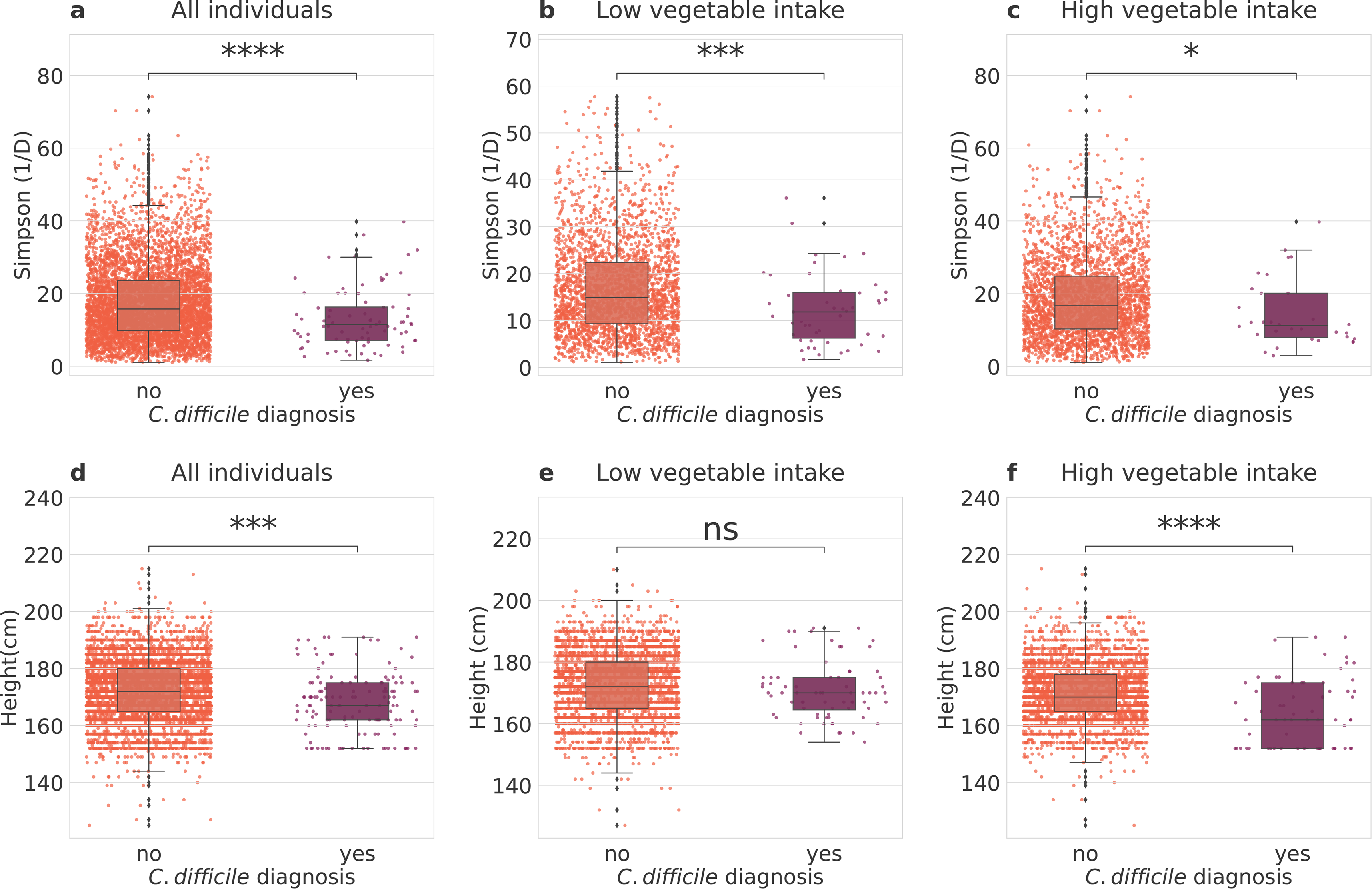
Exploring interactions between height, gut Simpson’s Diversity, diet, and self-reported history of CDI in the American Gut cohort. (A) Individuals who have reported a history of CDI have a significantly lower mean Simpson’s Diversity (n=9,113). (B) For individuals who reported low vegetable intake (n=4,383), mean Simpson’s Diversity was significantly lower in individuals who had a history of CDI. (C) Individuals who reported higher vegetable intake (n=4,730) also show significantly lower Simpson’s Diversity among those who reported a history of CDI. (D) The mean height of individuals who reported a history of CDI was significantly lower than individuals who did not. (E) There were no differences in height between people with and without a history of CDI if they had low vegetable intake. (F) Individuals with a history of CDI were significantly shorter than those who did not among individuals with high vegetable intake. Brackets with stars indicate various levels of significance when performing an independent t-test: **** signifies *p* ≤ 0.001, *** signifies 0.0001 < *p* ≤ 0.001, ** signifies 0.001 < *p* ≤ 0.01, and * signifies 0.01 < *p* ≤ 0.05.

We hypothesized that this association between height, diversity, and CDI could be influenced by diet, as prior research has shown that increasing consumption of a larger variety of plants is positively associated with gut microbiome alpha-diversity. We partitioned our analysis by self-reported vegetable consumption, looking at differences in height between those with or without a history of CDI in high and low vegetable consumption groups. Of the individuals who reported low vegetable intake,the mean height of individuals with and without a history of CDI was not significantly different (**Fig. 5E**, Welch’s t-test p=0.19). However, of the individuals who reported high vegetable intake, the mean heights of individuals who reported a history of CDI were significantly shorter than those who had not (**Fig. 5F**, *p* ≤ 0.0001). Gut alpha-diversity was always significantly lower in individuals with a history of CDI, independent of vegetable intake groups (**Fig 5B** and **5C**, *p* = 0.0018 and *p* = 0.028, for high and low intake, respectively).

We next ran binomial regression, correcting for a number of covariates, with CDI history as the dependent variable and height and vegetable consumption as independent variables. CDI history was significantly associated with vegetable consumption, even when correcting for age, sex, bowel movement frequency, and height (logistic regression β = -0.6682, *p* = 0.007), whereas height was not (*p* = 0.828). To follow up on this result, we ran a formal mediation analysis, with bootstrapping (simulations=5000). When we classified vegetable consumption as a treatment, Simpson’s Diversity as a mediator, and CDI history as an outcome, we found that the average causal mediated effect (ACME), the average direct effect (ADE), and total effect were all statistically significant (p=0.002, p=0.008, p=0.006, respectively), with average estimated coefficients of -0.001, -0.009, and -0.009, respectively (**Figure 6A**). 7.7% of the effect of vegetable consumption on CDI history were mediated through Simpson’s Diversity (**Figure 6A**; p=0.004). In the case of classifying height as a treatment, Simpson’s Diversity as a mediator, and CDI history as an outcome, the ACME was negative and statistically significant (ACME=-0.0004, p< 2E-16), but the Average Direct Effect (ADE) and the total effect were not statistically significant (**Figure 6B**, ADE=0.0008, p=0.73 and total effect = 0.0004, p=0.87). These results tentatively suggest that whatever effect that height may have on CDI history is completely mediated by Simpson’s Diversity.

**Figure 6.**
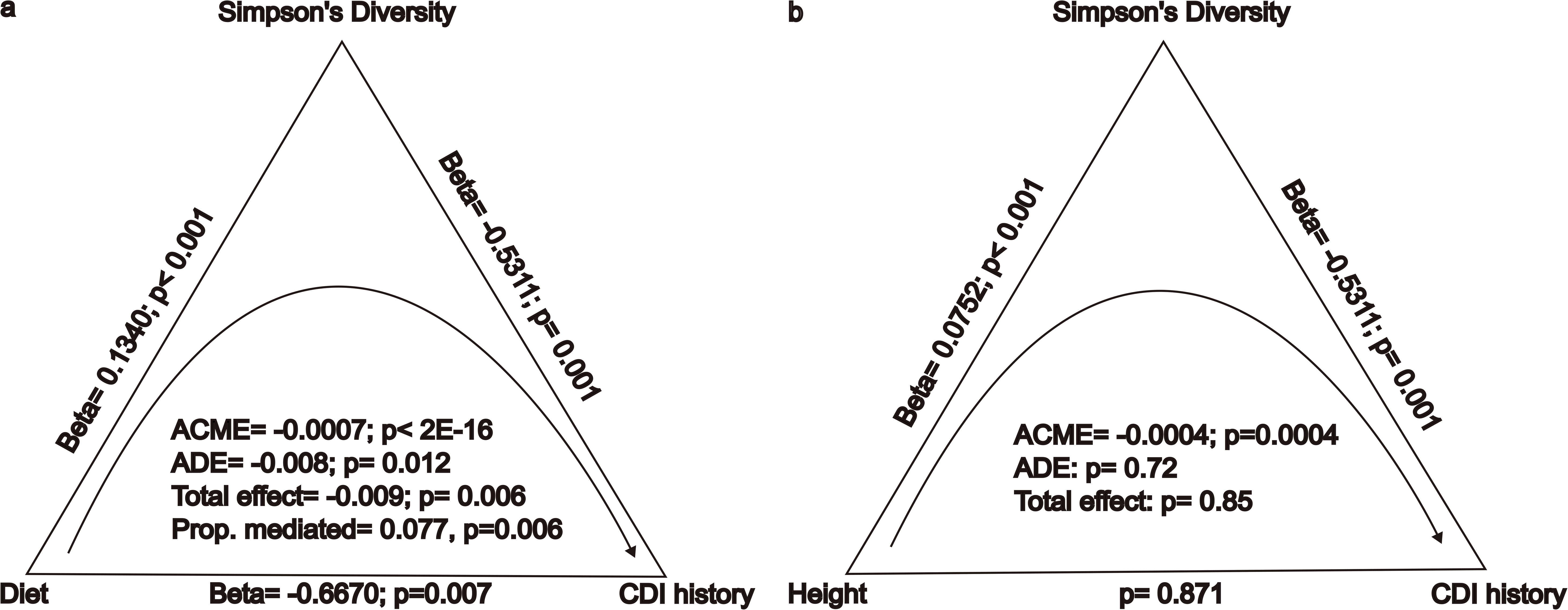
American Gut cohort mediation analysis investigating whether or not the effects of vegetable consumption and height on CDI history are mediated by Simpson’s Diversity. All regressions reported include the covariates: age, sex, BMI, and BMF. (A) A mediation analysis assigning vegetable consumption as a treatment, Simpson’s Diversity as a mediator, and CDI history as an outcome showed that the ACME, ADE, and total effect coefficients were negative and statistically significant (p<0.05). A diet higher in vegetable consumption was positively associated with Simpson’s Diversity (p<0.05). Both higher vegetable consumption and Simpson’s Diversity were significantly, negatively associated with reported CDI history (p<0.05). (B) A mediation analysis assigning height as a treatment, Simpson’s Diversity as a mediator, and reported CDI history as an outcome found that the ACME was the only statistically significant effect (p<0.05). Height was significantly, positively associated with Simpson’s Diversity, but not CDI history. Simpson’s Diversity was significantly negatively associated with CDI history (p<0.05).

## Discussion

Across three vertebrate cohorts and two large human cohorts with a rich set of relevant covariates, we demonstrated that there is a weak, but robust association between gut microbiome alpha-diversity and body size. We hypothesized that IBT provides a mechanistic explanation for this phenomenon, where larger guts are ecologically equivalent to larger islands. Indeed, we show how this pattern naturally emerges from an IBM designed to simulate IBT in a gut-like environment with unidirectional flow. Finally, we explore how the relationship between human height and gut alpha-diversity may be relevant to enteric bacterial infections, for which lower gut alpha-diversity is a known risk factor ^7,25,26^. We found that individuals who reported a history of CDI in the American Gut cohort were shorter than average and had lower gut alpha-diversity. Partitioning the analysis by vegetable consumption influenced this relationship, and a joint model that included both vegetable consumption and height showed how diet appears to be a much stronger driver of CDI risk than height.

### Bigger animals harbor more bacterial taxa in their guts

Consistent with prior literature, we found that vertebrate body size was positively associated with alpha-diversity ^12^. Across vertebrate data sets, body size explained approximately half of the variation in gut microbiome Simpson’s Diversity, for both legacy electrophoretic-based methods of detecting 16S phylotypes and for 16S amplicon sequencing (**Fig. 1**).

Literature on the human gut has reported many variables that can affect microbiome alpha-diversity ^8,16,27,28^. For example, prior work in less-than-healthy older individuals who live in assisted living facilities has shown a decline in alpha-diversity with age ^29^, while other studies in healthier older people and in community-dwelling centenarians have shown a decline in core taxa and increased alpha-diversity with age ^24,27,30^. In addition to age, sex has been associated with gut microbiome diversity, where females tend to have higher gut alpha-diversity than males ^27,31^. Obesity and BMI are negatively associated with gut alpha-diversity, likely due to lower dietary fiber intake and higher levels of systemic inflammation ^28,32^. Finally, BMF is negatively associated with gut alpha-diversity, with individuals experiencing constipation showing higher gut diversity and individuals experiencing diarrhea showing lower gut diversity ^33^. To complicate matters further, these demographic variables are highly interrelated. For example, females, on average, exhibit lower BMFs, shorter heights, and higher fruit and vegetable consumption than males, and the cumulative effects of these entanglements on diversity are difficult to predict ^27,33,34^. Fortunately, we were able to correct for all of these potentially confounding variables in our regressions, and we found that the association between height and gut alpha-diversity were robust to the inclusion of these other variables across two large, independent cohorts (**Table 1**).

### Application of the IBT to simulated guts of varying lengths

We hypothesized that these body-size versus gut microbiome alpha-diversity SARs could be explained by IBT, a simple neutral theory that shows how bigger islands harbor a larger number of ecologically-equivalent species through stochastic demographic processes like immigration/emigration and birth/death. We built a simple IBM, designed to approximate IBT in the gut, to show that gut length is indeed positively associated with species diversity (**Fig. 4**). In our model, we sampled from a mainland community with an exponential species abundance distribution (i.e., a small number of abundant species and a heavy-tail of low-abundance species). The amount of variance in Simpson’s Diversity explained by system length in the IBMs was on par with what we observed across both vertebrates and human body size scales (**Figs. 1-3**). Specifically, over a six orders of magnitude size range, we observe 43-58% of empirical variance in diversity explained across vertebrates and 34% of variance explained across IBM models. Over a smaller, two-fold, size range we saw 5-6% of empirical variance in diversity explained across humans and 7% of simulated variance explained across IBMs. The relationship between system length and diversity appeared to behave somewhat asymptotically in the vertebrate body size range (**Fig. 4A**), which was similar to what we and others have empirically observed (**Fig. 1A**) ^35^.

### Investigating the clinical implications of IBT in humans

An intact gut microbiota is a barrier against many infectious diseases ^19,20,36^. For example, CDI is the most common form of hospital-acquired colitis, and susceptibility to this disease is strongly related to common disruptions to the commensal gut microbiota, such as antibiotic treatment or diarrhea ^7,26,37,38^. CDI infections are initially treated with oral antibiotics, and recurrent illness is common, especially in those starting out with lower gut microbiome alpha-diversity ^7,25,39^. Thus, our results suggest that height may be a weak predictor of CDI susceptibility due to its influence on alpha-diversity. Indeed, we found that, on average, individuals reporting a history of CDI were shorter and had lower gut alpha-diversity than those who had no history of CDI (**Fig. 5**). However, this rather subtle association did not retain significance in a covariate-adjusted regression model.

Prior research, across both human populations and vertebrate species, has shown that gut alpha-diversity is higher in herbivores than in omnivores or carnivores, independent of the observed body-size and alpha-diversity scaling relationship ^8,12,40^. This can be attributed to the greater diversity of indigestible polysaccharides, like fibers and complex starches, in plants, which are fermented into organic acids by our commensal gut microbiota ^41^. We hypothesized that diet may be able to rescue the effect of height on gut alpha-diversity in shorter individuals, thus reducing their CDI risk. In order to test this hypothesis, we ran a formal mediation analysis using height or high vegetable consumption as the treatment, Simpson’s Diversity as the mediator, and CDI history as the outcome (**Figure 6**). We found that higher vegetable consumption was negatively associated with having a history of CDI, and that this effect was partially mediated by Simpson’s Diversity. When we defined height as the treatment, we found that the mediation effect (ACME) was statistically significant, but the direct and total effects were not. Classical mediation analysis does not consider the significance of the ACME in the absence of a significant direct or total effect ^42^. However, in certain scenarios, this conservative approach might miss true mediation effects, like when the treatment and mediator effects on the outcome are of opposite sign and together they cancel out the total effect. Alternatively, one can use bootstrapping to estimate confidence intervals for the indirect effect, as we did here, allowing for the estimation of ACME in the absence of direct or total effects ^43^. Thus, we show that height has a significant indirect effect on reported CDI history through its influence on Simpson’s Diversity, despite not having a detectable direct effect (**Fig. 6**). However, diet showed a much stronger association with both diversity and CDI history. This is good news for shorter individuals, suggesting that lifestyle interventions, like a higher-vegetable diet, can compensate for the negative influence of shorter stature on gut microbiome alpha-diversity.

In summary, we find a consistent association between body size and gut microbiome alpha-diversity across vertebrates and within human populations. The association between human height and gut microbiome alpha-diversity, while somewhat weak, was robust to the inclusion of several relevant covariates known to influence gut alpha-diversity, including age, sex, BMI, and BMF, across two large, independent cohorts. We showed that this macroecological scaling phenomenon could be explained by IBT, and that IBT simulations closely matched empirical observations. Finally, we explored how the relationship between human height and gut alpha-diversity is potentially relevant to CDI risk and how dietary patterns, like vegetable intake, may help mitigate this risk.

## Materials and Methods

### Published data sets

In order to investigate the relationship between vertebrate body size and gut microbiome alpha-diversity, we used three datasets from Godon et al. 2016, Song et al. 2020, and Groussin et al. 2017, respectively ^12,23,44^.

The Godon et al. 2016 dataset included pre-calculated Simpson’s Diversity ^12^, derived from single-strand conformation polymorphism (SSCP) capillary electrophoresis fluorescence patterns of amplicons from the V3 region of the 16S rRNA gene ^45^, and body mass for 71 vertebrate species, where bacterial diversity was assessed by extracting DNA from feces.

Song et al. 2020 included 16S rRNA amplicon sequencing data derived from 1,373 samples from 164 vertebrate species’ fecal samples. Samples that showed signs of contamination, from juveniles/newborn individuals, and from diseased individuals were removed. Moreover, duplicate samples from the same individual were removed and samples were included only if the host species had been sampled at least twice. ASVs that were not of bacterial origin were removed. Samples with no information on country of origin and/or information on the preservative used were removed as well.

Groussin et al. 2017 includes 16S amplicon sequencing data derived from fecal samples from 33 mammalian species. The OTU table was downloaded from Groussin’s publicly available Github repository ‘MammalianGuts’, and had been processed as described in Groussin et al. 2017 (https://github.com/mgroussi/MammalianGuts). Animal masses were not included from this study, but average masses for each species were manually curated from literature and from the AnimalTraits Database ^46–70^.

The data collected from the Arivale cohort included height, BMI, sex, BMF, and 16S amplicon data derived from stool samples. We used the mbtools workflow (https://github.com/gibbons-lab/mbtools) to denoise the 16S data. Moreover, training DADA2 error models and removal of chimeric reads using DADA2 was done separately for each sequencing run to generate sequence variants for each sample ^71^. The RDP classifier and the SILVA database (version 132) was used for taxonomy assignment, where species-level taxonomy were identified via exact alignment to SILVA reference sequences when possible ^72,73^. DECIPHER was used to align sequence variants ^74^. The multiple sequence alignment was trimmed by removing positions that consisted of more than 50% gaps, with a resulting core alignment of 420 base pairs.

We downloaded the American Gut data from figshare (https://doi.org/10.6084/m9.figshare.6137192.v1), with the metadata including self-reported height, weight, sex, age, BMF, CDI history, and vegetable intake frequency. We used the self-reported height and weight to calculate BMI for each participant. The sOTU table had been trimmed to a read length of 125 nucleotides and had been processed by Deblur^75^. Moreover, because samples had been sent via mail at room temperature, samples with obvious bacterial blooms that occurred in the sample tubes were removed. Samples with fewer than 1250 sequences were not included in the sOTU table. Subsequent filtering on the data was performed in order to avoid including erroneous self-reported data. We filtered out individuals who were less than 18 due to the confounding nature of age on gut microbiome diversity. Additionally, we removed samples from participants who reported a height greater than 244 centimeters or a height less than 122 centimeters, and we removed participants who reported a weight greater than or equal to 300 kilograms.

In order to investigate the relationship between human height and gut microbiome alpha-diversity, we used the Arivale cohort and the American Gut cohort. The Arivale and American Gut cohort metadata included information about each individual’s age, sex, height, BMI, and BMF. The way that BMF was recorded for the American Gut individuals was different from how it was recorded for the Arivale cohort individuals. In the Arivale cohort questionnaire data, participants were prompted to respond to : “I have bowel movements’’ with the options: “2 or fewer times per week”, “3-6 times per week”, “1-3 times daily”, and “4+ times daily”. The American Gut bowel questionnaire prompted individuals to respond to: “How many times do you have a bowel movement in an average day?”, where individuals could choose from the options: “Less than one”, “one”, “two”, “three”, “four”, or “five or more”. Because measures of BMF in each cohort were not directly comparable in each cohort, we standardized BMF, BMI, age, log-Simpson’s Diversity, and log-height by calculating their *Z*-score (*Z* = (*x* - mean(*x*))/std(*x*)) in each cohort prior to statistical analyses.

### Sequencing data processing and diversity calculations

We used Qiime 2-2022.8 on all ASV and OTU tables to determine the rarefaction depth, to rarefy our data, and to compute Simpson’s Diversity. Rarefaction depth was determined by sample frequency minimum, which was elucidated by Qiime’s summarize function. The sampling depth for each dataset was: Song et al. 2020 (sampling depth: 5,035), Groussin et al. 2017 (sampling depth: 1,266), Arivale cohort (sampling depth: 13,700), American Gut cohort (sampling depth: 1,250). Because Qiime 2-2022.8 returns Simpson’s Diversity as 1-D, we converted the Simpson’s Diversity returned by Qiime (Q_simpson) values to Simpson’s Diversity(1/D) using the numpy package and the formula 1/D = 1/(1-Q_simpson).

### Simulations

In our IBM, the gut is represented by a system in 1D space. Prior to each simulation, system length, mainland population abundance distribution parameters, reproductive rate, death rate, and immigration rate are randomly selected from defined ranges. The amount of simulations to run, and whether they are to be run on a large- or small-scale are selected by the user. When the system and its parameters have been initialized, individuals are generated and enter at one end of the system. Similar to a tube, individuals are able to enter the system at one end and exit from the other. Each time step of the simulation will track the flow of individuals within the system, removing individuals as they flow out, as well as generating new individuals or killing existing individuals within the system based on the reproduction and death rate. The immigration rate will define how many individuals will be added to the system per time step.

When running simulations at larger scales (spanning six orders of magnitude), a single entity in the model was coded as representing 100 individuals in order to increase memory efficiency. Subsequent diversity metrics were calculated by scaling the number of entities and the length of the system by a factor of 100.

### Statistical analyses

Ordinary Least Squares regression was used to test the association between vertebrate body size and gut microbiome alpha-diversity, using the following formula: log-Simpson’s Diversity ∼ log-mass(kg). We used the numpy package to log transform the Simpson’s Diversity (1/D) and the mass(kg) for all vertebrate datasets. The association between height and gut microbiome alpha-diversity in the Arivale and the American Gut cohorts was also tested using OLS regression, with the following formula (same covariates for each cohort): log-Simpson’s Diversity ∼ log-height + age + sex + BMI + BMF. In order to assess the significance of the OLS model, we used an ANOVA F-test to compare a reduced model with the formula: Simpson’s Diversity ∼ age + sex + BMI + BMF to the full model, which includes height, described above. We used Welch’s t-test in order to compare the mean heights of individuals with and without a history of CDI, both in the entire cohort, as well as within the groups ‘Low vegetable intake’ and ‘high vegetable intake’. We defined individuals with ‘high vegetable intake’ as individuals who ate vegetables at least once a day. Individuals who ate their vegetables less than once a day were placed in the ‘Low vegetable intake’ category. In order to test the relationships between height, vegetable consumption, and reported history of CDI, we used Generalized Linear Modeling with a binomial logit link using the formula: CDI history ∼ height + vegetable consumption + age + sex + BMI + BMF. We conducted our mediation analyses in R, using the “mediation” package ^76^, with height or diet as the treatment, alpha-diversity as the mediator, and CDI history as the response. The significance threshold for all tests was set at p<0.05.

## Data and code availability

All code, notebooks, and intermediate data files related to data analysis and IBM simulations can both be found in the following GitHub repository: https://github.com/Gibbons-Lab/IBT-and-the-Gut-Microbiome. Raw data from the Godon et al. 2016, Song et al. 2020, and Groussin et al. 2017 studies can be accessed in the original papers, or above in the ‘*Published data sets*’ section ^12,23,44^. Qualified researchers can access the full Arivale deidentified dataset supporting the findings in this study for research purposes through signing of a data use agreement. Enquiries to access the Arivale data can be made at and will be responded to within seven business days.

## Acknowledgements

We thank members of the Gibbons Lab for helpful feedback and suggestions on this work. We also thank Florent Mazel for providing the quality-filtered ASV tables and metadata for the Song et. al. dataset. This work was supported by a Washington Research Foundation Distinguished Investigator Award and by startup funds from the Institute for Systems Biology (to SMG). Research reported in this publication was also supported by the National Institute of Diabetes and Digestive and Kidney Diseases of the National Institutes of Health (NIH) under award no. R01DK133468 (to SMG.).

## Notes

https://github.com/Gibbons-Lab/IBT-and-the-Gut-Microbiome

### Competing Interest Statement

The authors have declared no competing interest.

## References

1. Martino, C. et al. Microbiota succession throughout life from the cradle to the grave. Nat. Rev. Microbiol. 20, 707–720 (2022).

2. Sender, R., Fuchs, S. & Milo, R. Revised Estimates for the Number of Human and Bacteria Cells in the Body. PLoS Biol. 14, e1002533 (2016).

3. Fan, Y. & Pedersen, O. Gut microbiota in human metabolic health and disease. Nat. Rev. Microbiol. 19, 55–71 (2021).

4. Diener, C. et al. Genome-microbiome interplay provides insight into the determinants of the human blood metabolome. Nat Metab 4, 1560–1572 (2022).

5. Cox, T. O., Lundgren, P., Nath, K. & Thaiss, C. A. Metabolic control by the microbiome. Genome Medicine 14, (2022).

6. Khan, I. et al. Mechanism of the Gut Microbiota Colonization Resistance and Enteric Pathogen Infection. Front. Cell. Infect. Microbiol. 11, 716299 (2021).

7. Pakpour, S. et al. Identifying predictive features of Clostridium difficile infection recurrence before, during, and after primary antibiotic treatment. Microbiome 5, 148 (2017).

8. McDonald, D. et al. American Gut: an Open Platform for Citizen Science Microbiome Research. mSystems 3, (2018).

9. Xu, Z. & Knight, R. Dietary effects on human gut microbiome diversity. Br. J. Nutr. 113 Suppl, S1–5 (2015).

10. Asnicar, F. et al. Blue poo: impact of gut transit time on the gut microbiome using a novel marker. Gut 70, 1665–1674 (2021).

11. Yassour, M. et al. Natural history of the infant gut microbiome and impact of antibiotic treatment on bacterial strain diversity and stability. Sci. Transl. Med. 8, 343ra81 (2016).

12. Godon, J.-J., Arulazhagan, P., Steyer, J.-P. & Hamelin, J. Vertebrate bacterial gut diversity: size also matters. BMC Ecol. 16, 12 (2016).

13. Li, S.-P. et al. Island biogeography of soil bacteria and fungi: similar patterns, but different mechanisms. ISME J. 14, 1886–1896 (2020).

14. Matthews, T. J., Triantis, K. A. & Whittaker, R. J. The Species–Area Relationship: Theory and Application. (Cambridge University Press, 2020).

15. Connor, E. F. & McCoy, E. D. The statistics and biology of the species-area relationship. Am. Nat. 113, 791–833 (1979).

16. Manor, O. et al. Health and disease markers correlate with gut microbiome composition across thousands of people. Nat. Commun. 11, 5206 (2020).

17. MacArthur, R. H. & Wilson, E. O. The Theory of Island Biogeography. (Princeton University Press, 2001).

18. van Werkhoven, C. H. et al. Incidence and predictive biomarkers of Clostridioides difficile infection in hospitalized patients receiving broad-spectrum antibiotics. Nat. Commun. 12, 2240 (2021).

19. Pereira, F. C. & Berry, D. Microbial nutrient niches in the gut. Environ. Microbiol. 19, 1366– 1378 (2017).

20. Kamada, N., Chen, G. Y., Inohara, N. & Núñez, G. Control of pathogens and pathobionts by the gut microbiota. Nat. Immunol. 14, 685–690 (2013).

21. Jernigan, A. & Hestekin, C. Capillary electrophoresis single-strand conformational polymorphisms as a method to differentiate algal species. J. Anal. Methods Chem. 2015, 272964 (2015).

22. Song, S. J. et al. Comparative Analyses of Vertebrate Gut Microbiomes Reveal Convergence between Birds and Bats. MBio 11, (2020).

23. Groussin, M. et al. Unraveling the processes shaping mammalian gut microbiomes over evolutionary time. Nat. Commun. 8, 14319 (2017).

24. Wilmanski, T. et al. Gut microbiome pattern reflects healthy ageing and predicts survival in humans. Nat Metab 3, 274–286 (2021).

25. Pérez-Cobas, A., Moya, A., Gosalbes, M. & Latorre, A. Colonization Resistance of the Gut Microbiota against Clostridium difficile. Antibiotics vol. 4 337–357 Preprint at https://doi.org/10.3390/antibiotics4030337 (2015).

26. VanInsberghe, D. et al. Diarrhoeal events can trigger long-term Clostridium difficile colonization with recurrent blooms. Nat Microbiol 5, 642–650 (2020).

27. de la Cuesta-Zuluaga, J. et al. Age- and Sex-Dependent Patterns of Gut Microbial Diversity in Human Adults. mSystems 4, (2019).

28. Wilmanski, T. et al. Blood metabolome predicts gut microbiome α-diversity in humans. Nat. Biotechnol. 37, 1217–1228 (2019).

29. Jeffery, I. B., Lynch, D. B. & O’Toole, P. W. Composition and temporal stability of the gut microbiota in older persons. ISME J. 10, 170–182 (2016).

30. Biagi, E. et al. Gut Microbiota and Extreme Longevity. Curr. Biol. 26, 1480–1485 (2016).

31. Wilmanski, T., Gibbons, S. M. & Price, N. D. Healthy aging and the human gut microbiome: why we cannot just turn back the clock. Nat Aging 2, 869–871 (2022).

32. Duan, M. et al. Characteristics of gut microbiota in people with obesity. PLoS One 16, e0255446 (2021).

33. Johnson, J. P. et al. Generally-healthy individuals with aberrant bowel movement frequencies show enrichment for microbially-derived blood metabolites associated with impaired kidney function. bioRxiv (2023) doi:10.1101/2023.03.04.531100.

34. Kim, Y. S., Unno, T., Kim, B. Y. & Park, M. S. Sex Differences in Gut Microbiota. World J. Mens Health 38, 48–60 (2020).

35. Matthews, T. J., Triantis, K. A. & Whittaker, R. J. The Species-Area Relationship: Theory and Application. (Cambridge University Press, 2021).

36. Honda, K. & Littman, D. R. The microbiome in infectious disease and inflammation. Annu. Rev. Immunol. 30, 759–795 (2012).

37. Tomkovich, S., et al. An Osmotic Laxative Renders Mice Susceptible to Prolonged Clostridioides difficile Colonization and Hinders Clearance. mSphere 6, e0062921 (2021).

38. Bignardi, G. E. Risk factors for Clostridium difficile infection. J. Hosp. Infect. 40, 1–15 (1998).

39. Johnson, S. Recurrent Clostridium difficile infection: a review of risk factors, treatments, and outcomes. J. Infect. 58, 403–410 (2009).

40. Losasso, C. et al. Assessing the Influence of Vegan, Vegetarian and Omnivore Oriented Westernized Dietary Styles on Human Gut Microbiota: A Cross Sectional Study. Front. Microbiol. 9, 317 (2018).

41. Cronin, P., Joyce, S. A., O’Toole, P. W. & O’Connor, E. M. Dietary Fibre Modulates the Gut Microbiota. Nutrients 13, (2021).

42. Baron, R. M. & Kenny, D. A. The moderator-mediator variable distinction in social psychological research: conceptual, strategic, and statistical considerations. J. Pers. Soc. Psychol. 51, 1173–1182 (1986).

43. Mackinnon, D. P., Lockwood, C. M. & Williams, J. Confidence Limits for the Indirect Effect: Distribution of the Product and Resampling Methods. Multivariate Behav. Res. 39, 99 (2004).

44. Mazel, F., Guisan, A. & Parfrey, L. W. Transmission mode and dispersal traits correlate with host specificity in mammalian gut microbes. Mol. Ecol. (2023) doi:10.1111/mec.16862.

45. Liu, Q., Scaringe, W. A. & Sommer, S. S. Discrete mobility of single-stranded DNA in non-denaturing gel electrophoresis. Nucleic Acids Res. 28, 940–943 (2000).

46. Herberstein, M. E. et al. AnimalTraits - a curated animal trait database for body mass, metabolic rate and brain size. Scientific Data 9, 1–11 (2022).

47. Laursen, L. & Bekoff, M. Loxodonta Africana. (1978).

48. Attias, N., Gurarie, E., Fagan, W. F. & Mourão, G. Ecology and social biology of the southern three-banded armadillo (Tolypeutes matacus; Cingulata: Chlamyphoridae). J. Mammal. 101, 1692–1705 (2020).

49. Fischer, J. et al. The Natural History of Model Organisms: Insights into the evolution of social systems and species from baboon studies. (2019) doi:10.7554/eLife.50989.

50. Réale, D., Festa-Bianchet, M. & Jorgenson, J. T. Heritability of body mass varies with age and season in wild bighorn sheep. Springer Nature 526–532 (1999).

51. Trani, M. K., Mark Ford, W. & Chapman, B. R. The Land Manager’s Guide to Mammals of the South. (2007).

52. Bauchot, R. & Stephan, H. DONNEES NOUVELLES SUR L’ENCEPHALISATION DES INSECTIVORES ET DES PROSIMIENS. 30, 160–196 (1966).

53. Kes Hillman-Smith, A. K. & Groves, C. P. Diceros Bicornis. (1994).

54. New data on the status and distribution of the bush dog (Speothos venaticus): Evaluating its quality of protection and directing research efforts. Biol. Conserv. 141, 2494–2505 (2008).

55. Basal rate of metabolism and temperature regulation in Goeldi’s monkey (Callimico goeldii). Comp. Biochem. Physiol. A Mol. Integr. Physiol. 135, 279–290 (2003).

56. Benatti, H. R., et al. Morphometric Patterns and Blood Biochemistry of Capybaras (Hydrochoerus hydrochaeris) from Human-Modified Landscapes and Natural Landscapes in Brazil. Veterinary Sciences 8, (2021).

57. Reamer, L. A. et al. Validation and Utility of a Body Condition Scoring System for Chimpanzees (Pan troglodytes). Am. J. Primatol. 82, e23188 (2020).

58. Boddy, A. M. et al. Comparative analysis of encephalization in mammals reveals relaxed constraints on anthropoid primate and cetacean brain scaling. J. Evol. Biol. 25, 981–994 (2012).

59. Dawson, T. J., Grant, T. R. & Fanning, D. Standard Metabolism of Monotremes and the Evolution of Homeothermy. Aust. J. Zool. 27, 511–515 (1979).

60. Metabolic rates of three gazelle species (Nanger soemmerringii, Gazella gazella, Gazella spekei) adapted to arid habitats. Mamm. Biol. 80, 390–394 (2015).

61. Crile, G. A Record of the Body Weight and Certain Organ and Gland Weights of 3690 Animals. (1940).

62. Bertelsen, M. F. Giraffidae. Fowler’s Zoo and Wild Animal Medicine, Volume 8 602 (2015).

63. Wilson, D. E. & Hanlon, E. Lemur catta (Primates: Lemuridae). mmsp 42, 58–74 (2010).

64. Sha, J. C. M. Comparative diet and nutrition of frugivorous and folivorous primates at the Singapore Zoo. JZAR 2, 54–61 (2014).

65. Gerstner, K., Liesegang, A., Hatt, J.-M., Clauss, M. & Galeffi, C. Seasonal body mass changes and feed intake in spectacled bears (Tremarctos ornatus) at Zurich Zoo. JZAR 4, 121–126 (2016).

66. Cain, J. W., Krausman, P. R. & Germaine, H. L. Antidorcas marsupialis. Mammalian Species 1–7 (2004).

67. Lurz, P. W. W., Fielding, I. & Hayssen, V. Callosciurus prevostii (Rodentia: Sciuridae). Mammalian Species 49, 40–50 (2017).

68. Hoefs, M. The thermoregulatory potential of Ovis horn cores. Can. J. Zool. 78, 1419–1426 (2000).

69. Liu, L. et al. The Visayan Warty Pig (Sus cebifrons) Genome Provides Insight Into Chromosome Evolution and Sensory Adaptation in Pigs. Mol. Biol. Evol. 39, (2022).

70. Saarinen, J., Cirilli, O., Strani, F., Meshida, K. & Bernor, R. L. Testing Equid Body Mass Estimate Equations on Modern Zebras—With Implications to Understanding the Relationship of Body Size, Diet, and Habitats of Equus in the Pleistocene of Europe. Front. Ecol. Evol. 9, 622412 (2021).

71. Callahan, B. J. et al. DADA2: High-resolution sample inference from Illumina amplicon data. Nat. Methods 13, 581–583 (2016).

72. Lan, Y., Wang, Q., Cole, J. R. & Rosen, G. L. Using the RDP classifier to predict taxonomic novelty and reduce the search space for finding novel organisms. PLoS One 7, e32491 (2012).

73. 25 years of serving the community with ribosomal RNA gene reference databases and tools. J. Biotechnol. 261, 169–176 (2017).

74. Wright, E. S. DECIPHER: harnessing local sequence context to improve protein multiple sequence alignment. BMC Bioinformatics 16, 322 (2015).

75. Amir, A. et al. Deblur Rapidly Resolves Single-Nucleotide Community Sequence Patterns. mSystems (2017) doi:10.1128/msystems.00191-16.

76. Tingley, D., Yamamoto, T., Hirose, K., Keele, L. & Imai, K. mediation: R Package for Causal Mediation Analysis. J. Stat. Softw. 59, 1–38 (2014).

77. Warnke, P. Mitteilung neuer Gehirn-und Körpergewichtsbestimmungen bei Saugern. J. Psychol. Neurol. 13, 355–403 (1908).

